# First direct observation of ‘elongated’ conformational states in α-synuclein upon liquid-liquid phase separation

**DOI:** 10.1101/2022.04.20.488894

**Authors:** Daniele Ubbiali, Marta Fratini, Lolita Piersimoni, Christian H. Ihling, Marc Kipping, Ingo Heilmann, Claudio Iacobucci, Andrea Sinz

## Abstract

α-Synuclein (α-syn) is an intrinsically disordered protein (IDP) that undergoes liquid-liquid phase separation (LLPS), fibrillation, and forms insoluble intracellular Lewy’s bodies in neurons, which are the hallmark of Parkinson’s Disease (PD). Neurotoxicity precedes the formation of aggregates and is probably related to LLPS of α-syn in the cell. The molecular mechanisms underlying the early stages of LLPS are still elusive. To obtain structural insights into α-syn upon LLPS, we take advantage of cross-linking/mass spectrometry (XL-MS) and introduce an innovative approach, termed COMPASS (COMPetitive PAiring StatisticS). COMPASS unravels transient interactions between α-syn molecules in liquid droplets. In this work, we show that the conformational ensemble of α-syn shifts from a ‘hairpin-like’ structure towards more ‘elongated’ conformational states upon LLPS. We obtain insights into the critical initial stages of PD and establish a novel mass spectrometry-based approach that will aid to solve open questions in LLPS structural biology.

## Introduction

Liquid-liquid phase separation (LLPS) of proteins has recently emerged as a highly relevant biological phenomenon that modulates several physiological and pathological processes^[1]^. The ability of forming phase-separated droplets is a characteristic of intrinsically disordered proteins (IDPs)^[2]^. IDPs lack a defined structure, but rather populate a wide ensemble of different conformational states^[3]^. The reason of this plasticity is related to the high number of states that minimize IDP inherent *frustration*^[4]^. Often, IDPs’ primary structure consists of a high number of charged amino acids over hydrophobic ones that guarantees a lack of a hydrophobic folded core^[5–7]^. Individual IDPs can establish transient and multivalent intra- and inter-molecular interactions, eventually leading to liquid condensate formation^[6]^. As proposed by Pappu et al., proteins that undergo LLPS possess adhesive domains, termed ‘stickers’, alternated by disordered and flexible regions, termed ‘spacers’. These structural elements can weakly and promiscuously interact with each other creating a hub of inter-protein interactions within the proteinaceous droplet. These interactions create a structurally heterogeneous higher order assembly^[2,8]^. A number of intra-protein interactions in the dilute phase become inter-protein interactions in the condensed phase, resulting in a shift of the conformational ensemble of an IDP upon LLPS^[9,10]^.

In cells, LLPS generates membrane-less organelles, covering a large number of compositions, structural features, and functions^[1]^. In a similar fashion, proteins that are prone to fibrillation, such as TDP-43^[11]^, FUS^[12]^, Tau^[13]^ and α-syn ^[14–16]^, are involved in e.g. amyotrophic lateral sclerosis (ALS), Alzheimer’s disease (AD), and Parkinson’s Disease (PD). They have been shown to undergo LLPS prior to their fibrillation and the formation of amyloid-like structures.

In the case of α-syn, the neurotoxicity that causes neuronal death in PD precedes the formation of α-syn fibrils and intracellular deposits of the protein, termed Lewy’s Bodies^[17]^. LLPS in α-syn might be the triggering factor leading ultimately to the development of PD. Therefore, the structural characterization of α-syn in the early stages of PD development is the key for developing novel therapeutic approaches in the future^[14]^.

α-Syn is a small (14.5 KDa) IDP that consists of two distinct regions, the membrane-binding region and the acidic region (Figure 1a). The amphipathic membrane-binding region contains seven repeat membrane binding motifs and includes the positively charged *N*-terminal region and the hydrophobic non-amyloid component (NAC) region. The *C*-terminal acidic region is thought to engage in long-range interactions with the *N-* terminal region of α-syn ^[15]^. This might result in the protection of the NAC region, thus autoinhibiting LLPS. Interestingly, high α-syn concentrations trigger the formation of liquid droplets (Figure 1b)^[15,16]^. During the early stages of LLPS, α-syn remains mainly monomeric^[14,18,19]^. The formation of oligomers and fibrils increases over time, suggesting that LLPS might be a critical step to nucleate α-syn aggregation^[14,15,20]^. NMR spectroscopy^[7]^, and low resolution methods like Förster Resonance Energy Transfer (FRET)^[21]^ and XL-MS^[22]^ can provide structural information within protein liquid droplets.

**Figure 1.**
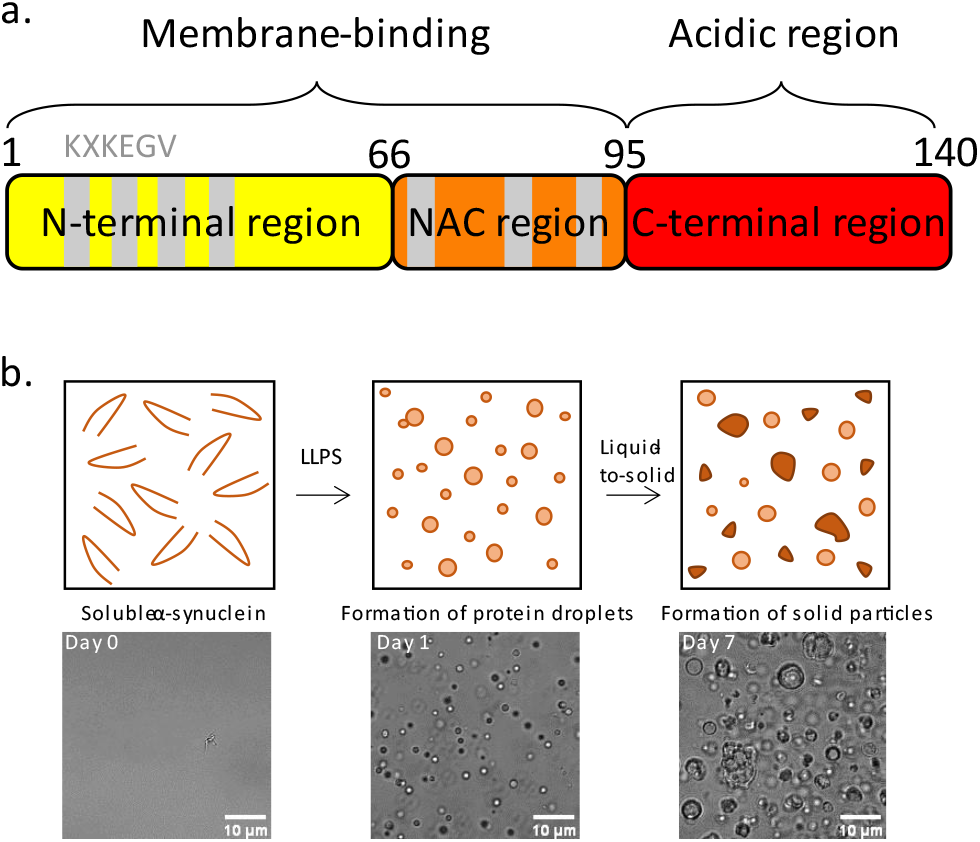
a) Schematic presentation of α-syn, N-terminal region (yellow), NAC region (orange), C-terminal acidic region (red), membrane binding motifs (grey). b) Time-dependent maturation of α-syn. At day 0 no LLPS is observed. At day 1 LLPS is visible. At day 7 aggregation occurs.

In this work, we further advanced the XL-MS approach for studying LLPS by introducing the innovative COMPASS method. XL-MS makes use of bifunctional reagents to covalently bridge side chains of amino acids that are in proximity in proteins, acting as “molecular rulers”^[23]^. We applied XL-MS to 1:1 mixtures of ^14^N- and ^15^N-labeled α-syn, while monitoring LLPS by microscopy. This allowed us distinguishing between intra- and inter-protein cross-links in α-syn that were then quantified with COMPASS. Inter-protein cross-links, triggered by transient protein-protein interactions, increase across the whole α-syn sequence. The distinct positions of these cross-links lead us to hypothesize that α-syn monomers populate more ‘elongated’ conformational states upon LLPS.

## Results and Discussion

α-syn is an intriguing protein system as monomers oligomerize and ultimately form amyloid fibrils^[17,20]^. The disease-relevant oligomeric states of α-syn are challenging to study from a structural perspective. XL-MS approaches fail to distinguish intra- from inter-protein crosslinks in oligomeric complexes precluding the investigation of α-syn homodimer interfaces. To overcome this limitation stable isotope–labeling of proteins is usually employed where labeled and unlabeled proteins are mixed at equimolar ratios prior to cross-linking. Inter-protein cross-links exhibit a characteristic pattern “quartet” of signals with identical intensities in the mass spectra, corresponding to the ^14^N^14^N-, ^14^N^15^N-, ^15^N^14^N-, and ^15^N^15^N-species (Figure 2)^[24,25]^. This isotope-mixing cross-linking approach is however not sufficient to fully solve the puzzle of oligomeric protein complexes. Only cross-links between ^14^N- and ^15^N-peptides can unambiguously be assigned as inter-protein cross-links for deriving protein interfaces. The majority of cross-links bridge two ^14^N- or two ^15^N-peptides and are therefore not applicable to derive structural information.

**Figure 2.**
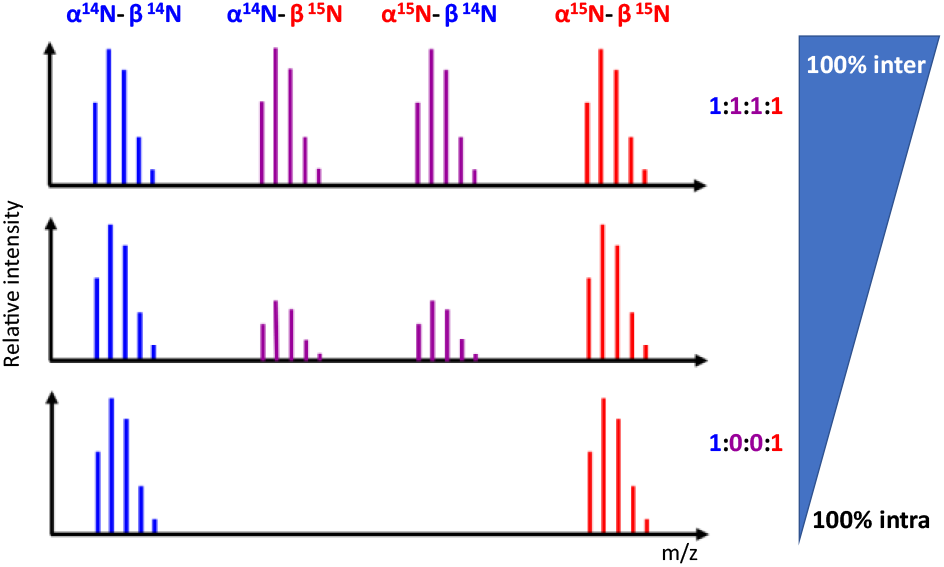
Schematic presentation of cross-link isotope patterns in mass spectra using 1:1 mixtures of ^14^N and ^15^N labeled proteins, depending on their intra-protein character. α refers to the heavier cross-linked peptide,. while β refers to the lighter one.

To fill this gap, we introduce an innovative strategy, termed COMPASS, which substantially extends the structural information that can be derived from XL-MS experiments.

The rationale behind COMPASS originates from the experimental observations that the four signals of inter-protein cross-links (^14^N^14^N, ^14^N^15^N, ^15^N^14^N, ^15^N^15^N) do not always show equal intensities in the mass spectra. Their intensities in the mass spectra might deviate from the statistical 1:1:1:1 ratio, with the external ^14^N^14^N, ^15^N^15^N species being more abundant (Figure 2). We interpreted this behavior as the result of a mixed inter- and intra-protein character of the respective cross-links. A cross-linker bridges two amino acid side chains within a protein monomer as well as between two monomers, depending on the relative spatial proximities of the respective residues in the protein assembly. As such, intra-protein cross-links will exclusively generate ^14^N^14^N ^15^N^15^N cross-links. The tendency of forming also inter-protein cross-links differs for every residue pair and results in the specific signal intensity ratios observed in the mass spectra (Figure 2). Based on the distinct ratios of the ^14^N^14^N, ^14^N^15^N, ^15^N^14^N, and ^15^N^15^N signals, COMPASS quantifies the percentage of the intra-protein character of a cross-link (*P_intra_* as:

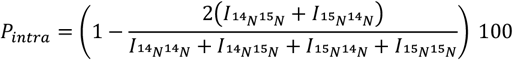

where *I*_14_*N*_14_*N*__, *I*_14_*N*_15_*N*__, *I*_15_*N*_14_*N*__, and *I*_15_*N*_15_*N*__, are the intensities of ^14^N^14^N, ^14^N^15^N, ^15^N^14^N, and ^15^N^15^N signals, respectively.

P_intra_ depends on the protein’s 3-D structure as well as the topology of the protein oligomer. A change of P_intra_ indicates a protein conformational change, e.g. upon LLPS or binding of a ligand.

Even proteins that do not form a well-defined structured oligomer, IDPs, and proteins undergoing LLPS can be investigated by the COMPASS approach. In fact, ubiquitous transient contacts in or between proteins can be stabilized by cross-linking, resulting in individual P_intra_ values < 100%. The higher the propensity of two amino acid side chains to form intra-molecular cross-links, the lower the probability to capture random inter-molecular contacts. Here we show the immense value of COMPASS to study the early events in LLPS of α-syn.

We incubated α-syn in a home-built moisture chamber (Figure S1) where the protein was aliquoted in a glass bottom multi-well plate. LLPS was observed over time by bright-field microscopy (Figure 1b). Cross-linking reactions were performed in triplicates using either disuccinimidyl dibutyric urea (DSBU) or 1-ethyl-3-(3-dimethylaminopropyl) carbodiimide hydrochloride/sulfo-N-hydroxysuccinimide (EDC/sulfo-NHS) cross-linkers. The complementarity of both cross-linking principles regarding their reactivities and spacer lengths (Figures S2 and S3) allowed targeting the complete sequence of α-syn. This is due to the peculiar clustering of primary amines (lysines) at α-syn’s N-terminus and carboxylic acid groups (aspartic and glutamic acids) at α-syn’s C-terminus. An initial cross-linking experiment was performed immediately after incubation (day 0), when the solution was clear (Figure 1b). LLPS of α-syn became visible after 24 h of incubation (day 1) (Figure 1b). Droplets presented a spherical morphology and were uniform in size (0.5-2 μm). We chose sampling of α-syn’s conformational ensemble at day 1 to focus on the early stages of LLPS, avoiding possible contaminations by α-syn aggregation. Our stringent approach allowed us to successfully monitor conformational changes in α-syn. We continued monitoring the LLPS and α-syn aggregation for one week (day 7) until the morphology and the size of all droplets became irregular, suggesting further phase transitions (Figure 1b).

Cross-linked α-syn samples from day 0 and day 1 were denatured with urea, fractionated by size exclusion chromatography (SEC), and analyzed by electrospray ionization mass spectrometry (ESI-MS); for details see Supporting Information. We identified cross-linked monomeric and dimeric α-syn alongside with residual unreacted protein (Figure S4). As expected, the majority of α-syn remained monomeric upon LLPS without any alteration of the monomer-to-dimer ratio (Figure S4). We successfully assigned cross-linked species containing different numbers of cross-linker molecules (Figures S5 and S6). In particular, the presence of α-syn dimers containing at least two cross-links indicate the coexistence of intra- and inter-protein cross-links, enabling the application of COMPASS (Figure S7).

High-resolution tandem mass spectra (MS/MS) were recorded and analyzed by the MeroX software^[26]^ to identify cross-linking sites. Signal intensities of ^14^N^14^N, ^15^N^14^N, ^14^N^15^N, and ^15^N^15^N cross-links in the mass spectra were extracted, and their intra-protein character (P_intra_) was calculated by applying Equation 1. We considered only cross-links (12 DSBU and 9 EDC/sulfo-NHS) that were accurately quantifiable in all three replicates of day 0 and day 1.

All cross-links together with their COMPASS quantitation are presented in Figure 3, while DSBU and EDC/sulfo-NHS data are presented separately.

**Figure 3.**
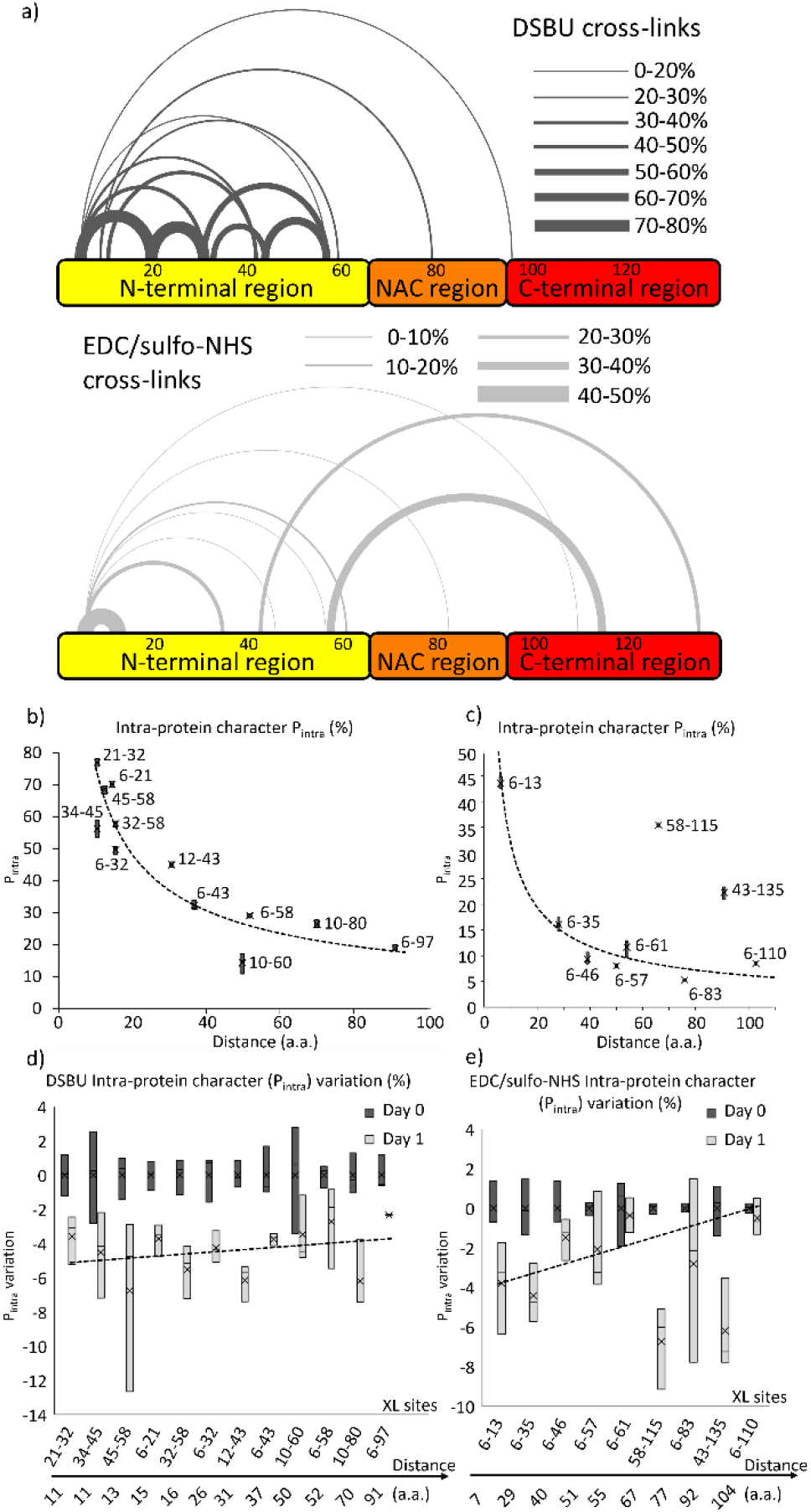
COMPASS reveals structural details of α-syn during early LLPS events. a) Quantitative visualization of the inter-protein character (P_intra_) of DSBU (dark gray) and EDC/sulfo-NHS (light gray) cross-links at day 0. Long-range cross-links in α-syn, involving the C-terminal region, with high inter-protein character defines the ensemble of “hairpin-like” structures. Boxplots of the P_intra_ of DSBU (b)) and EDC/sulfo-NHS (c)) cross-links at day 0 are plotted against the distances of the connected residues in the amino acid sequence. P_intra_ values are always >0 and exponentially decrease with the residue distance. This reveals an overall ‘elongated’, yet disordered, structure for the N-terminal and NAC regions in α-syn. The two EDC/sulfo-NHS cross-links involving the C-terminal region do not follow this P_intra_ trend of an overall ‘elongated’ structure. Boxplots of P_intra_ variations of each DSBU (d)) and EDC/sulfo-NHS (e)) cross-links at day 0 are reported. The average values of P_intra_ at day 0 are considered as references for each cross-link. All P_intra_ values decrease at day 1. The P_intra_ decrease becomes less substantial for long-range cross-links, which are characterized by a low P_intra_ at day 0. Long-range cross-links involving the C-terminal region of α-syn cluster apart from this distribution. Their P_intra_ decrease at day 1 suggest an ‘elongation’ of the auto-inhibited conformation upon LLPS.

It is worth noting that the absolute intra-protein characters of DSBU and EDC/sulfo-NHS cross-links cannot be directly compared as both cross-linking systems possess different spacer lengths and reactivities (Figure S4, S5). In this study, we focus on evaluating the intra-protein character of cross-links to study conformational changes in α-syn monomers upon LLPS. At day 0, before LLPS, DSBU cross-links involve residues of the N-terminal region and the NAC region (Figure 3a). The intra-protein character of cross-links in α-syn decreases with the distances in the primary structure (Figures 3a and 3b). Among the observed DSBU cross-links, Lys-21 and Lys-32 have the shortest distance (10 amino acids), and the highest intra-protein character, ≈77%. On the other hand, Lys-6 and Lys-97 have the longest distance (90 amino acids) and the lowest intra-protein character, ≈19%.

This suggests an overall extended structure of the N-terminal region and the NAC region in α-syn. Even residues that are far apart in α-syn’s amino acid sequence exhibit P_intra_ values > 0%. On the other hand, α-Syn also populates more compact structures within its conformational ensemble. P_intra_ values > 0% for DSBU cross-links implies that the N-terminal and NAC regions also contact other α-syn molecules. This is in agreement with the lack of a structurally well-defined α-syn dimer.

Interestingly, EDC/sulfo-NHS cross-links are distributed across the α-syn sequence (Figure 3a). The intra-protein character of EDC/sulfo-NHS cross-links involving the *N*-terminal region and the NAC region exhibit the same trends as observed for DSBU (Figures 3b and 3c). The P_intra_ values drop more rapidly with increasing distance in the amino acid sequence.

This behavior is due to the shorter distance of EDC/sulfo-NHS zero length cross-linker compared to DSBU (maximum Cα-Cα distance ~ 30Å).

On the other hand, EDC/sulfo-NHS cross-links between the acidic C-terminal region and the NAC region display a higher intra-protein character, even between residues that are far apart in the amino acid sequence (Figure 3a and 3c). This behavior can be explained as a preference of α-syn’s C-terminal region to populate conformations that directly contact the NAC region. Our cross-linking data can be best explained by the formation of a ‘hairpin-like’ structure in α-syn at day 0 (Figure 4).

**Figure 4.**
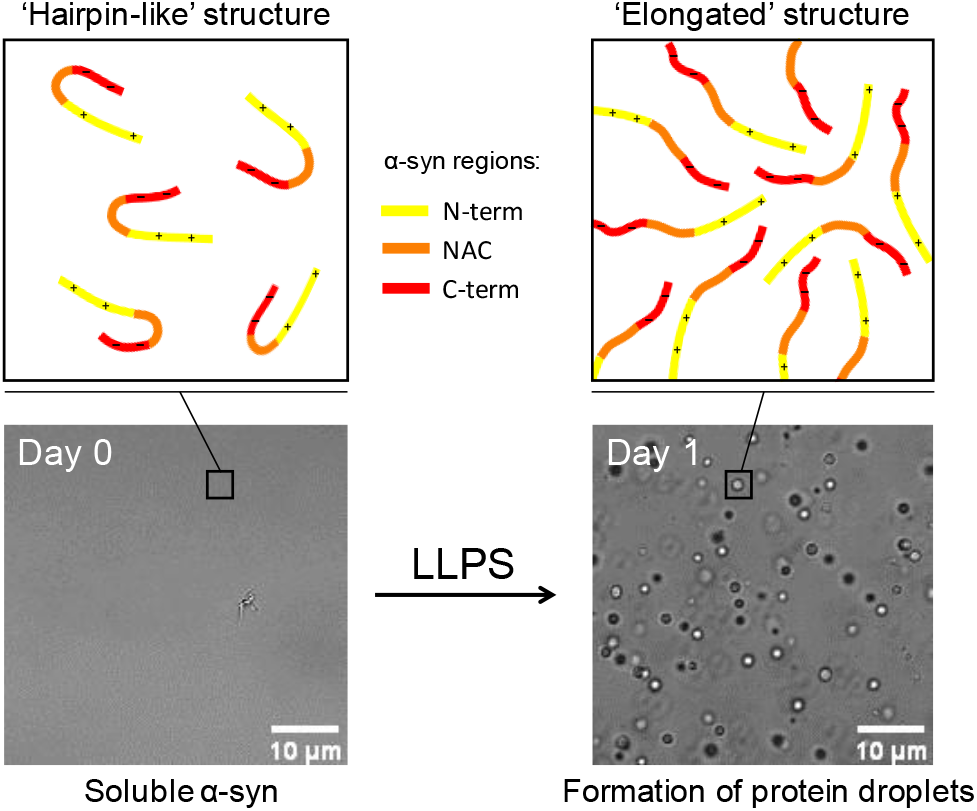
Cartoon representation of α-syn ‘hairpin-like’ and ‘elongated’ structures. In the ‘hairpin-like’ structure the positively charged N-terminal region engages in long-range interactions with the negatively charged C-terminal region, thus shielding the NAC domain. The ‘elongated’ structure of α-syn promotes inter-protein contacts and LLPS.

Recently, Sawner *et al.* observed that high salt concentrations promote LLPS of α-syn^[15]^. They speculated that charge neutralization, induced by increasing ionic strength at the positively charged N-terminal and the negatively charged C-terminal regions, modulates α-syn LLPS. This charge neutralization would reduce intramolecular interactions within α-syn.

Our data constitute the first direct observation of a long-range interaction at the molecular level between the C-and the N-terminal regions of α-syn prior to LLPS, which might form a ‘hairpin-like’ structure (Figure 4). This finding prompted us to investigate the P_intra_ values of cross-links identified at day 1 in more detail. Eventual Local variations of P_intra_ values will help in understanding the molecular mechanisms driving LLPS in α-syn. At day 1, P_intra_ of all DSBU and EDC/sulfo-NHS cross-links decrease (Figure 3d and 3e). As LLPS progresses, the local α-syn concentration increases and transient interactions between α-syn molecules become more frequent. DSBU captures a constant increase of those interactions in the N-terminal and NAC regions (Figure 3d). On the other hand, EDC/sulfo-NHS reveals a more pronounced increase of inter-protein cross-links involving the acidic C-terminal region compared to the other regions (Figure 3e). We rationalize this behavior as a shift of the conformational ensemble of α-syn from a ‘hairpin-like’ structure towards more ‘elongated’ conformations. The soluble ‘hairpin-like’ structure of α-syn at day 0 ‘elongates’ upon LLPS, enabling novel inter-protein interactions in the liquid condensates. This is the first observation at the molecular level to corroborate the hypothesis of an alteration of an initial auto-inhibited state of α-syn upon LLPS^[15]^.

## Conclusions

We successfully established and applied an innovative approach (COMPASS) for interpreting XL-MS data of α-syn, which can generally be used for investigating protein oligomers and IDPs. For the first time, we have obtained direct structural information about α-syn upon LLPS. α-Syn is present in solution as a monomer. The conformational ensemble of α-syn is in agreement with a ‘hairpin-like’ structure prior to LLPS (day 0). In particular, the C-terminal region of α-syn engages long-range interactions with the N-terminal region. The resulting shielding of the NAC region and sequestration of multivalent inter-protein interaction sites has already been hypothesized by Sawner *et al.* as inhibiting factor of α-syn LLPS^[15]^. Our data, obtained at the molecular level, are in agreement with that previous hypothesis. Upon LLPS, the ensemble of α-syn’s conformations seem to relax into more ‘elongated’ conformations. Elongation of the C-terminal region makes the membrane binding region of α-syn accessible to inter-protein interactions. While the conformational ensemble of α-syn shifts towards more ‘elongated’ conformations, α-syn itself remains monomeric, highly flexible and disordered upon LLPS. These molecular insights might form the basis for developing novel therapeutic strategies for treating PD.

## Experimental

Experimental procedures are provided in detail in the Supporting Information. MS data have been deposited to the ProteomeXchange Consortium via the PRIDE partner repository with the project accession PXD033205, username; password: LUQYIOsm

Expression and purification of α-syn were performed according to a previously described method^[27]^. α-syn was incubated in a moisture chamber for up to 7 days. Microscopy experiments were conducted using a Zeiss LSM880, with a 63X oil immersion objective, using the ZEN Black image analysis software. A ACQUITY UPLC Protein BEH SEC column (200Å, 1.7μm, 4.6mm x 300mm 10-500KDa (Waters) was employed in SEC experiments. A triple quadrupole XEVO TQ mass spectrometer (Waters), operating in full MS1 mode, was used in combination with SEC fractionation. SEC fractionation was performed manually. XL-MS experiments were performed with DSBU and EDC/sulfo-NHS at RT. Samples were then analyzed by LC/MS using a timsTOF Pro mass spectrometer (Bruker Daltonik). MS/MS spectra were annotated by the MeroX software and precursor intensities of cross-linked species were manually extracted.

## Acknowledgements

AS acknowledges financial support by the DFG (RTG 2467, project number 391498659 “Intrinsically Disordered Proteins – Molecular Principles, Cellular Functions, and Diseases”, INST 271/404-1 FUGG, INST 271/405-1 FUGG, and CRC 1423, project number 421152132), the Federal Ministry for Economic Affairs and Energy (BMWi, ZIM project KK5096401SK0), the region of Saxony-Anhalt, and the Martin Luther University Halle-Wittenberg (Center for Structural Mass Spectrometry). The authors are indebted to Prof. Tim Bartels for providing pTSara-NatB plasmid.

