## Supplemental_Materials for "First direct observation of ‘elongated’ conformational states in α-synuclein upon liquid-liquid phase separation"

### Table of Content

**Page Number**

**Materials and Methods ……………………………………………………………………………………………………3**

- Protein Expression and Purification
- Liquid-Liquid Phase Separation and Microscopy
- DSBU Cross-linking
- EDC Cross-linking
- SEC-ESI-MS and Fractionation
- Enzymatic Digestion
- Nano-HPLC-MS/MS
- MS Data Analysis
- COMPASS

**Amino Acid Sequence ……………………………………………………………………………………………..5**

**Additional Figures and Table …………………………………………………………………………………….6**

**Literature References ……………………………………………………………………………………………..14**

#### Materials and Methods

##### Protein Expression and Purification

Recombinant production of N-terminally acetylated α-synuclein (α-syn) was performed by co-expression of α-syn with N-terminal acetylase B (NatB)[1]. The α-syn gene was ligated into a pET21a(+) vector exploiting the XhoI and NdeI restriction sites. The pTSAra-NatB plasmid was kindly provided by Tim Bartels (Harvard Medical School) for co-expression. *Escherichia coli* BL21(DE3) cells were transformed with both plasmids. Cell growth and expression steps were performed as described previously[1], protein purification was done according to [2]. The protein was finally eluted with 20mM phosphate buffer (pH 7.4) and used for further analysis.

##### Liquid-Liquid Phase Separation and Microscopy

Protein concentrations were measured using a Nanodrop (Thermo Fisher Scientific) UV photometer (extinction coefficient at 280 nm: 5960 $M^{-1} \cdot cm^{-1}$, molecular weight: 14.5 kDa). $^{14}$N-α-syn and $^{15}$N-α-syn were 1:1 mixed (final concentrations of 100 µM each) in 20 mM phosphate buffer (pH 7.4). 100 µL of protein solution was placed in a µ-Slide 18-well glass bottom multi-well plate (Ibidi) for imaging and cross-linking reactions. The α-syn solution was incubated in a home-built moisture chamber (Figure S1). Bright field images were acquired with a Zeiss LSM880, with a 63X oil immersion objective using the ZEN Black image analysis software. The images were further processed with Fiji[3].

##### DSBU Cross-linking

DSBU (CF Plus Chemicals, 10 mM stock solution in DMSO) was added to α-syn to a final concentration of 200 µM. The mixtures were incubated for 60 minutes at room temperature. Cross-linked samples were stored at -80°C

##### EDC/sulfo-NHS Cross-linking

EDC and sulfo-NHS (25 mM stock solution in DMSO each) were added to α-syn to a final concentration of 500 µM. The mixtures were incubated for 90 minutes a room temperature. Cross-linked samples where stored at -80°C.

##### SEC-ESI-MS and SEC Fractionation

Cross-linked samples were dried in a vacuum concentrator and resuspended in 50 µl of 8 M urea, 400 mM ammonium bicarbonate. Reconstituted cross-linked samples were split in two aliquots of 10 µL and 40 µL, respectively.

The 10-µL aliquots of reconstituted cross-linked samples were analyzed by LC-MS/MS using an ACQUITY UPLC H-Class System PLUS (Waters) coupled to a XEVO TQ-XS mass spectrometer (Waters), equipped with Z-spray ion source (Waters). A SEC column (ACQUITY UPLC Protein BEH SEC, 4.6 mm x 300 mm, 200 Å, 1.7 µm, 10-500 kDa) was employed. Samples were eluted at 300 µL/min with 0.4% formic acid (Figure S4). The settings of the XEVO TQ-XS mass spectrometer were set as follows: Capillary voltage, 2 kV; sample cone voltage, 45 V; desolvation temperature, 400°C. MS data were acquired in full scan mode (*m/z* range 600-2000).

The remaining 40-µL aliquots of reconstituted cross-linked samples were manually fractionated to collect the α-syn dimer (retention times between 6 to 6.8 min) (Figure S4).

#### Enzymatic Digestion

All α-syn dimer fractions were dried in a vacuum concentrator. Reduction and alkylation steps were omitted due to the absence of cysteine residues in the amino acid sequence of α-syn. The samples were resuspended in 5 µl of 8 M urea in 400 mM ammonium bicarbonate. After 5 min of sonication, samples were diluted to 0.8 M urea, and incubated overnight at 37°C with 50 ng GluC (NEB). Subsequently, 50 ng trypsin (Gold, Promega) were added and incubated for 4 h at 37°C. All samples were acidified with TFA.

#### Nano-HPLC-MS/MS

Digested samples were analyzed by nano-HPLC-MS/MS on an UltiMate 3000 RSLC nano-HPLC system (Thermo Fisher Scientific) coupled to a timsTOF Pro mass spectrometer equipped with CaptiveSpray source (Bruker Daltonik). Peptides were trapped on a C18 precolumn (Acclaim PepMap 100, 300 µm × 5 mm, 5 µm, 100 Å, Thermo Fisher Scientific) and separated on a µPAC 50-cm column (PharmaFluidics). After trapping, peptides were eluted by a concave 90-min gradient from 3% (v/v) to 35% (v/v) ACN. For elution, a flow rate gradient was employed ranging from 900 to 600 nl/min in 15 min, followed by a constant flow rate of 600 nl/min. All LC separations were performed at room temperature.

For the settings of the timsTOF Pro mass spectrometer, the following parameters were adapted, starting with the parallel accumulation serial fragmentation (PASEF) method for standard proteomics. The values for mobility-dependent collision energy ramping were set to 95 eV at an inversed reduced mobility ($1/k_0$) of 1.6 V s/cm$^2$ and 23 eV at 0.73 V s/cm$^2$. Collision energies were linearly interpolated between these two $1/k_0$ values and kept constant above or below. No merging of TIMS scans was performed. Target intensity per individual PASEF precursor was set to 20,000. The scan range was set between 0.6 and 1.6 V s/cm$^2$ with a ramp time of 166 ms. 14 PASEF MS/MS scans were triggered per cycle (2.57 s) with a maximum of seven precursors per mobilogram. Precursor ions in the *m/z* range between 100 and 1700 with charge states ≥2+ and ≤8+ were selected for fragmentation. Active exclusion was enabled for 0.4 min (mass width 0.015 Th, 1/k0 width 0.015 V s/cm$^2$).

#### MS Data Analysis

Identification of cross-links was performed with MeroX 2.0.1.7. The following settings were applied: Proteolytic cleavage: *C*-terminal at Lys and Arg for (3 missed cleavages were allowed) and *C*-terminal at Asp and Glu (3 missed cleavages were allowed); peptide lengths of 5 to 30 amino acids; modifications: alkylation of Cys by iodoacetamide (fixed), oxidation of Met (variable); cross-linker specificity: Lys, Ser, Thr, Tyr, *N*-terminus (DSBU), Lys, N-terminus, Asp, Glu, *C*-terminus (EDC/sulfo-NHS); Lys, *N*-terminus, Asp, Glu, C-terminus (sulfo-SDA); search algorithm: RISEUP mode with up to three missing ions mode with minimum peptide score of 0; precursor mass accuracy: 10 ppm; fragment ion mass accuracy: 20 ppm; signal-to-noise ratio > 1.5; precursor mass correction enabled; 10% intensity as prescore cut-off; 1% false discovery rate (FDR) cut-off, and minimum score cut-off: 50. Mass recalibration was performed for all files.

MS data have been deposited to the ProteomeXchange Consortium via the PRIDE partner repository with the project accession PXD033205, username:; password: LUQYIOsm.

**Amino Acid Sequence of Human α-Synuclein**

MDVFMKGLSKAKEGVVAAAEKTKQGVAEAAGKTKEGVLYVGSKTKEGVVHGVATVAEKTK

EQVTNVGGAVVTGVTAVAQKTVEGAGSIAAATGFVKKDQLGKNEEGAPQEGILEDMPVDP

DNEAYEMPSEEGYQDYEPEA

**Additional Figures and Table**

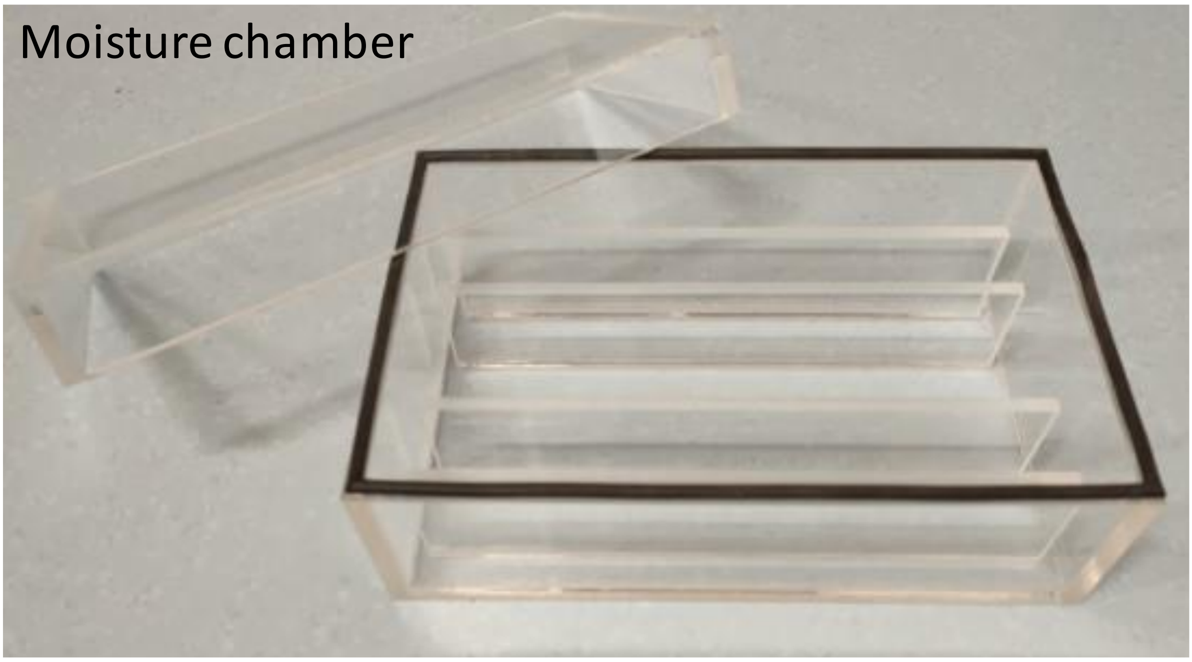


**Figure S1**. Home-built moisture chamber.

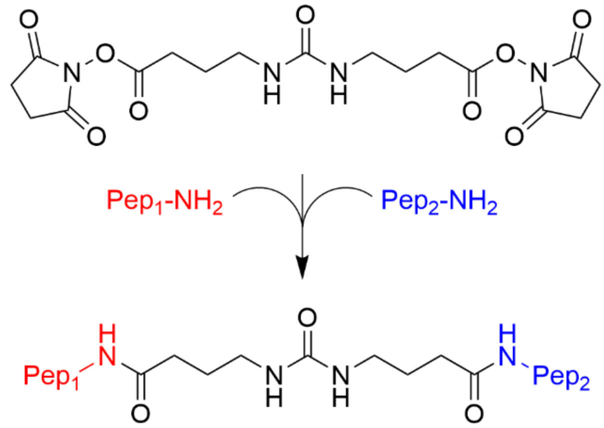


**Figure S2**. Reaction mechanism of DSBU cross-linker.

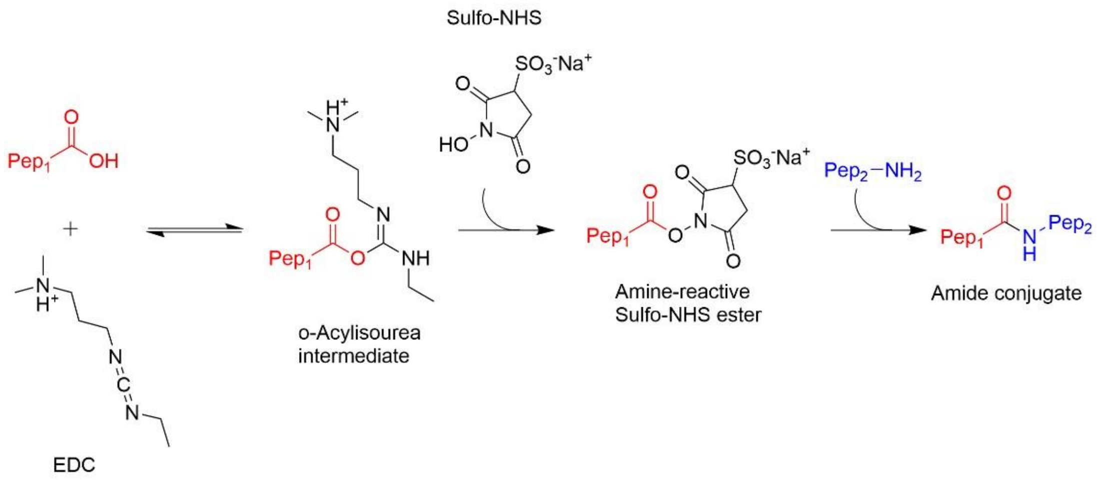


**Figure S3**. Reaction mechanism of EDC/sulfo-NHS cross-linker.

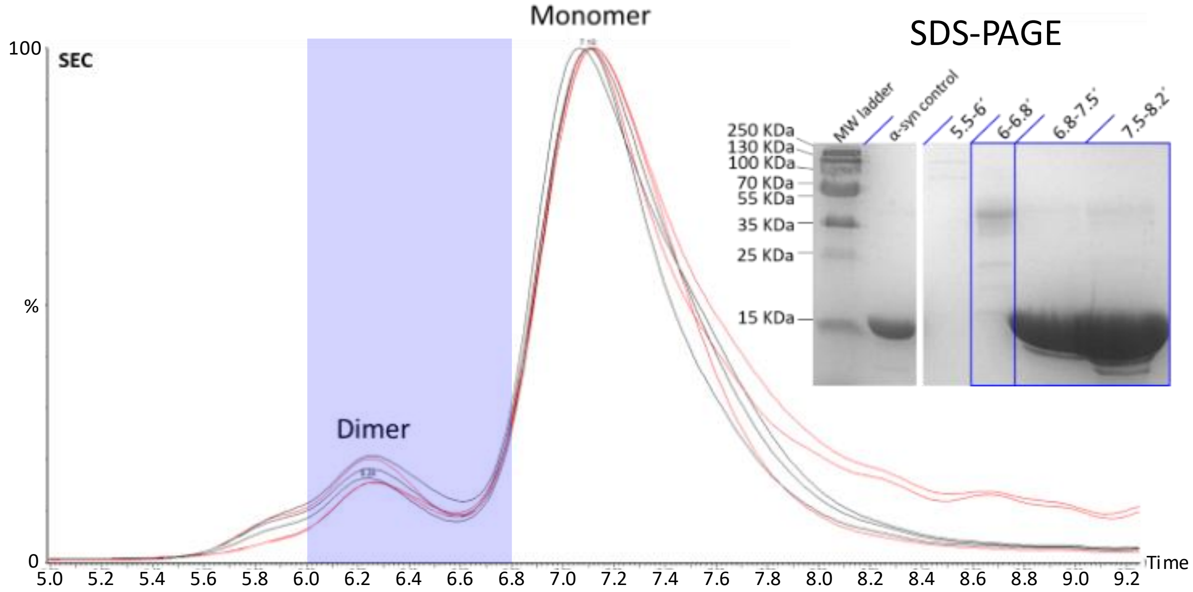


**Figure S4**. SEC-ESI-MS chromatograms of cross-linked α-syn samples. Total ion chromatogram (TIC) traces of day 0 sample triplicates, black solid lines; TIC traces of day 1 sample triplicates, red solid lines. SDS-PAGE analysis of collected SEC fractions is reported.

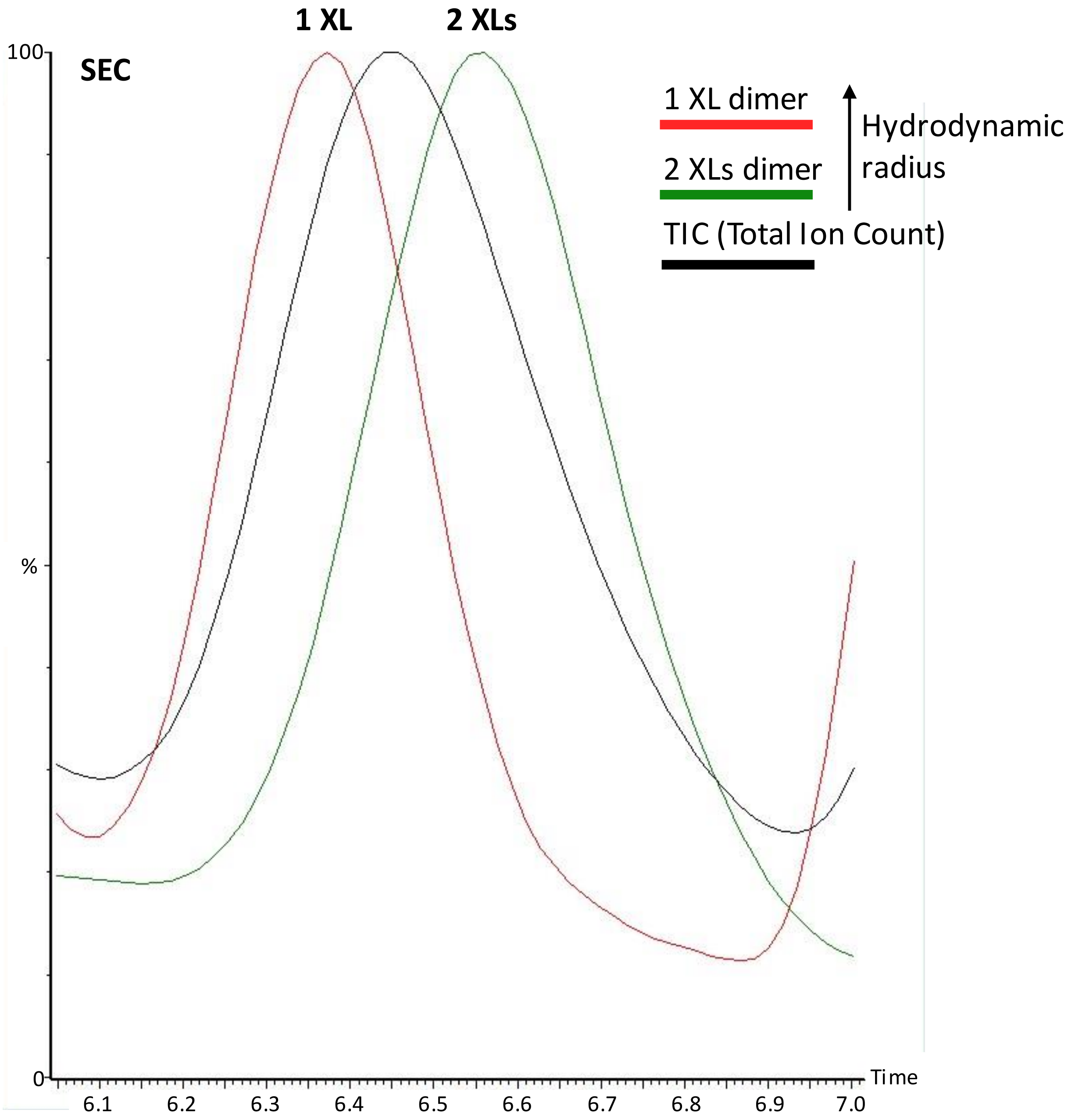


**Figure S5**. Chromatograms (SEC) of a cross-linked α-syn sample. TIC, solid black line; extracted ion chromatogram (XIC) of 1328.0-1328.4 *m/z* range corresponding to α-syn dimer with one DSBU modification, solid red line; XIC of 1360.4-1363.9 *m/z* range corresponding to of α-syn dimer with two DSBU modification, solid green line.

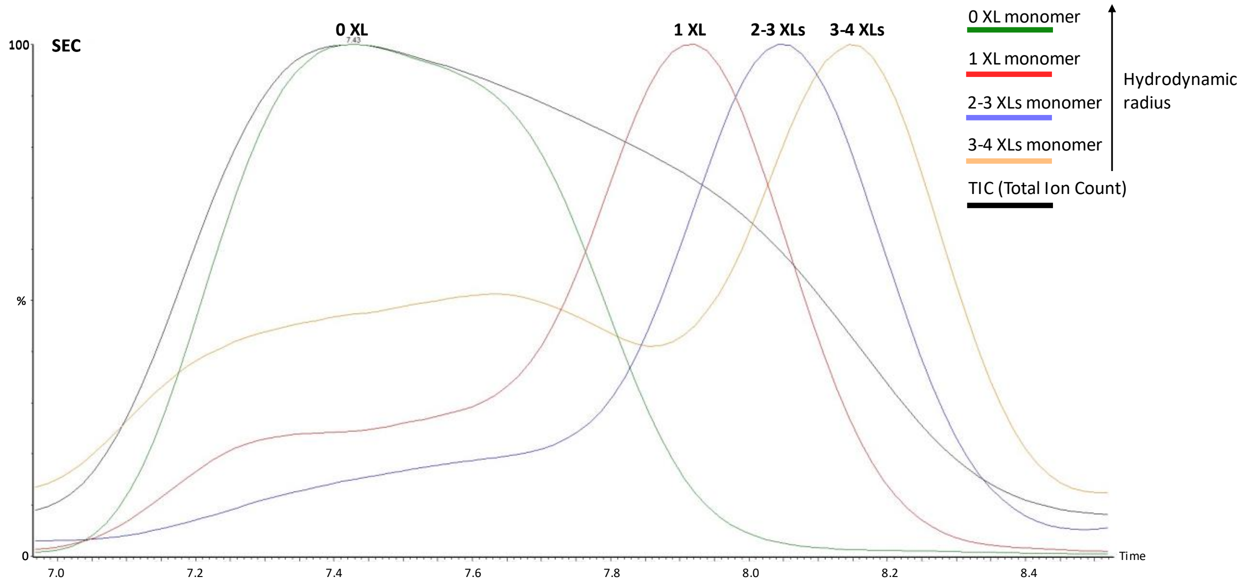


**Figure S6**. SEC-ESI-MS chromatograms of a cross-linked α-syn sample. TIC, solid black line; XIC of 1115.9-1116.9 *m/z* range corresponding to unmodified α-syn monomer, solid green line; XIC of 1131.2-1131.9 *m/z* range corresponding to α-syn monomer with one DSBU modification, solid red line; XIC of 1158.1-1162.0 *m/z* range corresponding to of α-syn monomer with two and three DSBU modifications, solid blue line; XIC of 1173.6-1178.0 *m/z* range corresponding to of α-syn monomer containing three and four DSBU modifications, solid orange line.

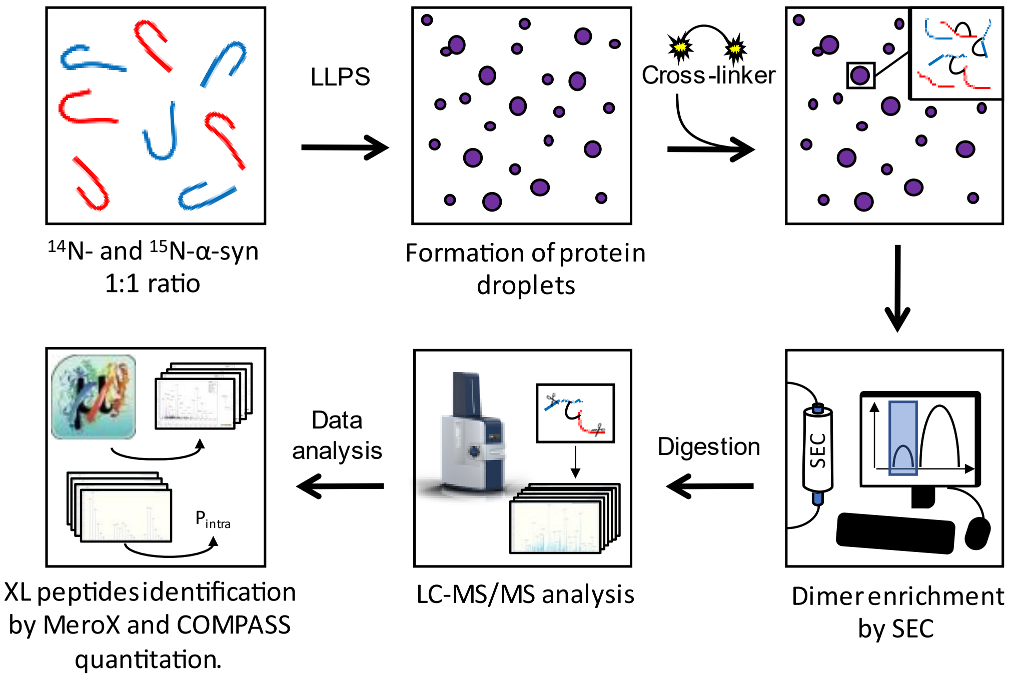


**Figure S7**. COMPASS workflow.

**Table S1.** Summary of DSBU and EDC/sulfo-NHS cross-links with $P_{intra}$ values (%) at day 0 and day 1.

| Cross-linker | Site1 | Site2 | Day0_1 (%) | Day0_2 (%) | Day0_3 (%) | Day1_1 (%) | Day1_2 (%) | Day1_3 (%) |
|---|---|---|---|---|---|---|---|---|
| DSBU | 34 | 45 | 53.4 | 56.4 | 58.7 | 49.0 | 54.0 | 52.0 |
| DSBU | 21 | 32 | 75.7 | 77.0 | 78.1 | 71.7 | 74.5 | 73.9 |
| DSBU | 45 | 58 | 66.9 | 69.3 | 68.7 | 65.4 | 55.7 | 63.6 |
| DSBU | 6 | 21 | 69.1 | 70.8 | 70.0 | 65.2 | 66.5 | 67.0 |
| DSBU | 32 | 58 | 56.6 | 58.6 | 58.1 | 50.5 | 53.6 | 52.6 |
| DSBU | 6 | 32 | 48.0 | 50.5 | 50.3 | 44.5 | 45.3 | 46.4 |
| DSBU | 12 | 43 | 45.7 | 44.7 | 44.2 | 39.2 | 37.5 | 39.5 |
| DSBU | 6 | 43 | 31.1 | 31.4 | 33.8 | 28.0 | 28.7 | 28.5 |
| DSBU | 6 | 58 | 28.3 | 29.6 | 29.3 | 27.2 | 28.3 | 23.6 |
| DSBU | 10 | 60 | 10.9 | 17.1 | 14.9 | 9.8 | 13.1 | 9.5 |
| DSBU | 10 | 80 | 25.3 | 26.1 | 27.6 | 18.9 | 22.6 | 18.9 |
| DSBU | 6 | 97 | 18.2 | 19.9 | 18.1 | 16.4 | 16.3 | 16.4 |
| EDC/sulfo-NHS | 6 | 13 | 43.4 | 45.5 | 43.5 | 42.4 | 40.9 | 37.8 |
| EDC/sulfo-NHS | 6 | 35 | 17.8 | 16.2 | 15.0 | 13.6 | 11.6 | 10.6 |
| EDC/sulfo-NHS | 6 | 46 | 8.8 | 10.9 | 8.8 | 8.9 | 6.9 | 8.3 |
| EDC/sulfo-NHS | 6 | 57 | 7.7 | 8.4 | 8.2 | 9.0 | 4.3 | 4.9 |
| EDC/sulfo-NHS | 6 | 61 | 12.4 | 9.9 | 13.1 | 10.6 | 11.4 | 12.3 |
| EDC/sulfo-NHS | 58 | 115 | 36.1 | 36.0 | 35.6 | 30.9 | 26.8 | 29.9 |
| EDC/sulfo-NHS | 6 | 83 | 5.3 | 5.1 | 5.6 | 6.8 | -2.5 | 3.2 |
| EDC/sulfo-NHS | 43 | 135 | 23.7 | 22.9 | 21.2 | 14.8 | 19.0 | 15.4 |
| EDC/sulfo-NHS | 6 | 110 | 8.4 | 8.6 | 8.8 | 9.1 | 7.3 | 7.9 |

#### Literature References


[1] M. Rovere, A. E. Powers, D. S. Patel, T. Bartels, *PLOS ONE* **2018**, *13*, e0198715.
[2] A. E. Powers, D. S. Patel, *Methods Mol Biol* **2019**, *1948*, 261–269.
[3] J. Schindelin, I. Arganda-Carreras, E. Frise, V. Kaynig, M. Longair, T. Pietzsch, S. Preibisch, C. Rueden, S. Saalfeld, B. Schmid, J.-Y. Tinevez, D. J. White, V. Hartenstein, K. Eliceiri, P. Tomancak, A. Cardona, *Nat Methods* **2012**, *9*, 676–682.
